# Influence of the male germline on B chromosome transmission in *D. melanogaster*

**DOI:** 10.64898/2026.10.08.756769

**Authors:** Natalie Warsinger-Pepe, Govind M Raman, Stacey L Hanlon

## Abstract

Selfish genetic elements that bias their inheritance to the detriment to their host are in an evolutionary arms race with host-derived suppressors that neutralize the spread of these elements. The supernumerary B chromosomes recently found in a mutant *Drosophila melanogaster* laboratory stock are subject to drive (>50% transmission) when passed through the female germline, but through the male germline they experience the opposite of drive, a phenomenon referred to as drag (<50% transmission). Here we show that males from this mutant stock do not have fertility defects, excluding the possibility that sperm carrying more B chromosomes are inviable. Though we revealed irregularities in the germline stem cell (GSC) niche, we show that other aspects of GSCs activity is normal: there is no delay in their mitoses and the B chromosomes appear to be symmetrically segregated during the GSC divisions, suggesting that the asymmetry of these divisions is not likely the source of B chromosome drag. Together, this work lays the foundation for future studies into the mechanism of paternal B chromosome drag in *D. melanogaster*, which is poised to significantly advance on our understanding of how various genetic and developmental contexts promote disparate chromosome behavior.

## INTRODUCTION

To achieve meiotic parity, chromosomes must segregate properly and without bias during the meiotic divisions. This faithful and equal transmission of chromosomes to the next generation results in an inheritance frequency that is Mendelian (50%), however selfish genetic elements can increase their transmission frequency to result in higher than Mendelian transmission, otherwise known as drive (Hanlon and Larracuente 2022). Nonessential, supernumerary chromosomes, termed B chromosomes (opposed to their essential counterparts, the “A” chromosomes), are selfish genetic elements that often exhibit drive during meiosis (Houben 2017; Warsinger-Pepe and Hanlon 2026). Recently, B chromosomes were identified in a mutant *Drosophila melanogaster* laboratory stock and were shown to be subject to meiotic drive when transmitted through the female germline (Hanlon and Hawley 2023). This *D. melanogaster* stock carries a null allele of *matrimony (mtrm)*, a female-specific meiotic gene, and the highly inverted balancer chromosome (*TM3)*, both of which were found to be necessary for strong B chromosome drive through the female germline (*yw;mtrm^126^/TM3,Sb,Ser;sv^spa-pol^* plus B chromosomes, herein referred to as *mtrm/TM3+*B). Though *mtrm* is only expressed in the female germline (Xiang et al. 2007), the B chromosomes are subject to reduced transmission (<50%), or drag, when transmitted through the male germline in the *mtrm/TM3+*B genetic background but not in a wild-type genetic background (*yw;sv^spa-pol^* plus B chromosomes, herein referred to as WT*+*B) that was generated after outcrossing the original stock to a lab wild-type stock (Hanlon and Hawley 2023). These opposing, non-Mendelian inheritance patterns indicate that a combination of the genetic background and sex-specific germline environment critically influence B chromosome heredity, but the mechanism(s) of how these contexts exert their influence is unknown.

The ability of this B chromosome to undergo drive in one context and drag in another is not entirely unexpected. Many selfish genetic elements are detrimental to their host (reviewed in (Warsinger-Pepe and Hanlon 2026)), creating an intragenomic conflict. One way to resolve this conflict is through the emergence of a host-derived genetic variant that opposes the proliferation of a selfish genetic element, triggering an evolutionary arms race that can lead to Red Queen dynamics (Levine 2026). Consistent with this model, several examples of suppressors that limit or eliminate the biased transmission of their respective drivers have been identified. For example, the *Segregation Distorter (SD)* drive system in *D. melanogaster* kills sperm that are packaged with its target, an *SD*-sensitive *Responder* (*Rsp*) locus (Larracuente and Presgraves 2012). Sperm that carry *SD* are unaffected because of the tight linkage of *SD* to a resistant variant of *Rsp*, resulting in the biased transmission of *SD* to progeny at the cost of reduced male fertility. As a result, natural populations of *D. melanogaster* have evolved suppressors that oppose *SD* activity and thus limit its spread (Chang et al. 2026). For B chromosomes, however, there are very few that have been shown to be capable of both drive and drag. Of these, the underlying biology responsible for their contrary segregation behavior is largely unexplored, making the B chromosomes in *D. melanogaster* an attractive model system for investigating how genetic and germline contexts influence chromosome segregation.

To initiate our investigation into how the male germline promotes B chromosome drag, we aimed to identify male germline phenotypes unique to the *mtrm/TM3+*B genetic background. We identified variable, but not reduced, fertility of *mtrm/TM3+*B males, indicating that the source of drag is not the loss or inviability of sperm with higher B chromosome copy numbers. We then focused on the beginning of spermatogenesis by examining the germline stem cell (GSC) niche and found irregular phenotypes that are unique to the *mtrm/TM3*+B genotype. Following this result, we analyzed the GSC divisions in several genetic backgrounds and conclude that there is no significant difference in their mitotic index. Due to the inherent asymmetry of GSC divisions, we also assessed the partitioning of the *AAGAT* satellite repeat that is enriched on the B chromosomes and found that its distribution between a GSC and its daughter gonialblast (GB) cell appears to be equal. By eliminating two common sources of transmission distortion (sperm inviability and transmission variance during the asymmetric GSC division), our study has made significant progress towards elucidating the mechanism of B chromosome drag in the *D. melanogaster* male germline.

## MATERIALS AND METHODS

### Drosophila husbandry and genetics

Unless otherwise noted, all stocks were reared at 25°C in a 12-hour light-dark cycling incubator on Nutri-Fly Bloomington Formulation Pre-Mixed Fly Food (Genessee Scientific #66-112), with propionic acid (64mM final concentration) and Tegosept (0.1% final concentration) and supplemented with active dry yeast. Fly sorting and collections were done using CO2 flowing from a pad. *Y w/Y,y^+^*;*sv^spa-pol^ +Bs* (*WT+B*) males were generated through a two-step crossing scheme. The first step was performed by crossing virgin/un-mated *y w/Y,y^+^;mtrm^126^/TM3,Sb,Ser;sv^spa-pol^ +Bs (mtrm/TM3+B)* females and *y w/Y,y^+^*;*sv^spa-pol^* (*WT*) males. *TM3,Sb Ser;sv^spa-pol^/+* progeny from the first step (virgin/unmated females and sibling males) were crossed to each other for the second step. *TM3,Sb Ser;sv^spa-pol^* -negative progeny (predicted to carry B chromosomes) were collected and aged as needed. *WT+B* larvae were generated by crossing *TM3,Sb Ser;sv^spa-pol^* -negative progeny (virgin/unmated females and sibling males) obtained from the second step cross. The full genotypes and their short names of all stocks used in this manuscript are listed in Supplemental Table 1.

### Sperm exhaustion assays

Sperm exhaustion assays were performed as described in (Chang et al. 2023) with slight modifications. Single, 2-5 day old virgin/un-mated males were paired in vials with ten, 2-5 day old, virgin/un-mated *y w* females using a mouth aspirator (no CO2) and reared at 25°C. Mating pairs were transferred to new vials (designated as a “Brood”) every 3-5 days for 13-14 days. P0 from the final vial were discarded 3-5 days later. All F1 were scored from each vial (Brood 1 through 4) until 17 days after initial pairing.

### Testes dissection and immunofluorescence

Protocol was modified from (Yamashita et al. 2003). Testes were dissected in 1X PBS and fixed in 1mL of 4% formaldehyde in 1X PBS in an Eppendorf tube on a nutator for 30 minutes at room temperature (RT). Fixed testes were then washed in 1X PBST (1X PBS containing 0.1% Triton X-100) for at least 2 hours at RT (or washed for 30 minutes at RT followed by an overnight incubation in 1X PBST at 4°C). Samples were then incubated with primary antibodies diluted in 1X PBST containing 3% bovine serum albumin (BSA) at 4°C overnight. Samples were washed three times in 1X PBST for 20-30 minutes each wash and then incubated with secondary antibodies in 1X PBST with 3% BSA at either RT in the dark for 2 hours or at 4°C overnight in the dark. If necessary, samples were washed as described above and then incubated a third time with mouse anti α-tubulin (IgG) conjugated to FITC (Sigma F2168, 1:250) in 1X PBST with 3% BSA at either RT in the dark for 2 hours or at 4°C overnight in the dark. Samples were washed once with 1X PBST and DAPI (4′,6-diamidino-2-phenylindole) (1:100) for 30 minutes at RT in the dark. Samples were washed two times in 1X PBST for 20 minutes each at RT in the dark. Samples were mounted in Prolong Glass Antifade mounting media (Fisher P36980) and stored coverslip down at 4°C overnight prior to imaging. Primary antibodies used were rat anti-vasa (DSHB: AB 760351, 1:300), mouse anti-Fasciclin III (gift from Mayu Inaba, DSHB: AB 528238, 1:20), and mouse anti-α-spectrin (DSHB: AB 528473, 1:50). Secondary antibodies used were Goat anti-Rat Alexa Fluor 546 (Thermo-Fisher A11081, 1:200-1:250) and Donkey anti-Mouse IgG Alexa Fluor 647 (Southern Biotech 6411-31, 1:250).

### Combined immunofluorescence and DNA fluorescent in situ hybridization (IF-FISH)

IF-FISH was performed as previously described (Meng and Yamashita 2025) with minor modifications. Testes were dissected and immunofluorescence was performed as described above. During the immunofluorescence protocol, EDTA was spiked into every solution to a final concentration of 1mM EDTA. Following the PBST washes after secondary antibody incubation, samples were postfixed in 4% paraformaldehyde (in 1X PBS, 1mM EDTA) for 30 minutes at room temperature (RT) in the dark. All subsequent steps were performed in the dark. Samples were washed in 1X PBST+1mM EDTA for 30 minutes. Fixed samples were incubated with 100μL RNase A solution (Sigma #10109142001, 2mg/mL in 1X PBST, no EDTA) at 37°C for 10 minutes. Samples were washed with 1X PBST+1mM EDTA for 5-10 minutes. Samples were then rinsed in 2X SSC-Tween20 (saline sodium citrate buffer with 0.1% Tween 20) +1mM EDTA. Samples were then washed at RT in 100μL of 2X SSC-Tween20 with increasing percentages of formamide three times for 15 minutes each wash (20%, 40%, then 50% formamide). Samples were further washed in 100μL of 2X SSC-Tween20 with 50% formamide for 30 minutes. Hybridization mix was added to each sample (40μL per sample, 45% formamide, 1mM EDTA, 2.5μM *AAGAT* conjugated to Alexa 488, 1.8X SSC, 9% dextran sulfate), gently mixed with a cut p200 pipette tip, and denatured at 91°C for 3 minutes. Samples were then wrapped in parafilm and aluminum foil and incubated overnight in a 37°C water bath. Samples were washed one time with 2X SSC-Tween20+1mM EDTA with DAPI (4′,6-diamidino-2-phenylindole, 1:100) for 30 minutes at RT. Samples were subsequently washed two times in 2X SSC-Tween20+1mM EDTA at RT for 20 minutes each wash. Samples were mounted in Prolong Glass Antifade mount (Fisher P36980) and stored coverslip down at 4°C overnight prior to imaging.

### Measuring relative corrected total fluorescence

Image quantification parameters were modeled from (Watase et al. 2022) with minor modifications. Prior to image acquisition for each apical tip, the signal intensity for *AAGAT* was manually adjusted to ensure no signal saturation. Confocal z-stacks were then obtained through the entire thickness of the apical tip of every testis analyzed. All images were processed and analyzed using Fiji. Usable pairs of germline stem cells (GSCs) and their associated daughter cell (gonial blast, GB) were identified if 1) the GSC was appropriately adjacent to the hub and with cytoplasmic vasa signal, 2) a fusome, visualized by α-spectrin, connected the cytoplasm of the GSC and an adjacent GB, and 3) the *AAGAT* signal for both GSC and GB existed in overlapping z-planes. For these GSC-GB pairs, all relevant slices were processed through Z-Project to Sum all values into one image. *AAGAT* signal per GSC or GB were manually circled using the Freehand Selection Tool to create a region of interest (ROI) to measure the area, mean value, and integrated density for each cell nucleus. Care was taken to avoid *AAGAT* signal from adjacent nuclei. A small portion of the nucleus within the ROI was separately circled to calculate the background levels in the *AAGAT* channel. The corrected total fluorescence (CTF) was calculated for each GSC and GB by CTF = Integrated Density – (Area*Mean of background reading). The GSC:GB ratio was calculated by dividing the GSC CTF by the GB CTF.

### Confocal fluorescent microscopy

Whole testes were imaged on an inverted Leica SP8 Spectral Confocal microscope housed by the Advanced Light Microscopy Facility at the University of Connecticut. Images were taken with a 63x oil immersion objective [numerical aperture (NA) = 1.40], processed using Fiji.

### Statistical analysis, graphical data representation, and figure assembly

GraphPad Prism was used for statistical analysis and graphical data presentation. Figures were created using Inkscape and Microsoft PowerPoint.

## RESULTS

### Sperm exhaustion assays reveal variable, but not reduced, male fertility

We first sought to identify any broad disruptions to fertility of *mtrm/TM3+B* males that may indicate areas of interest for disruptions in B chromosome transmission. To identify any subtle differences in fertility, sperm exhaustion assays were performed with *y w/Yy^+^* and *WT+B* control males to compare to males from the *mtrm/TM3+B* stock. We found no statistically significant differences in total progeny sired when comparing the three male genotypes assessed, which may be due to the large spread in progeny production for *mtrm/TM3+B* and *WT+B* males (Figure 1A). The sperm exhaustion assay also revealed no significant differences in the sex of the progeny sired by these three male genotypes (Supplemental Figure 1A), however *mtrm/TM3+B* males sired significantly more progeny carrying the third chromosome with the *mtrm* allele (61.88%) than progeny carrying the *TM3* balancer chromosome (38.12%) (Supplemental Figure 1B). We next measured the production of *X/X/Y* or *X/O* progeny in our sperm exhaustion assay to determine if nondisjunction (NDJ) was elevated in any of our genotypes and found no significant differences, indicating that chromosome segregation is not globally disrupted in our genotypes (Supplemental Figure 1C). It is interesting to note that *mtrm/TM3+B* males and *WT+B* males had higher median values of NDJ compared to *y w/Yy^+^* males (0.81%, 1.47%, and 0%, respectively) and that the majority of NDJ products were observed as *X/O* males (*mtrm/TM3+B* = 0.61%, *WT+B* = 1.46%) (Supplemental Figure 1C). Though we cannot ascertain if the NDJ event occurred in the female or male germline, we attribute the elevated NDJ frequency to the male genotype because the female genotype (*y w*) was held constant (see Methods).

**Figure 1:**
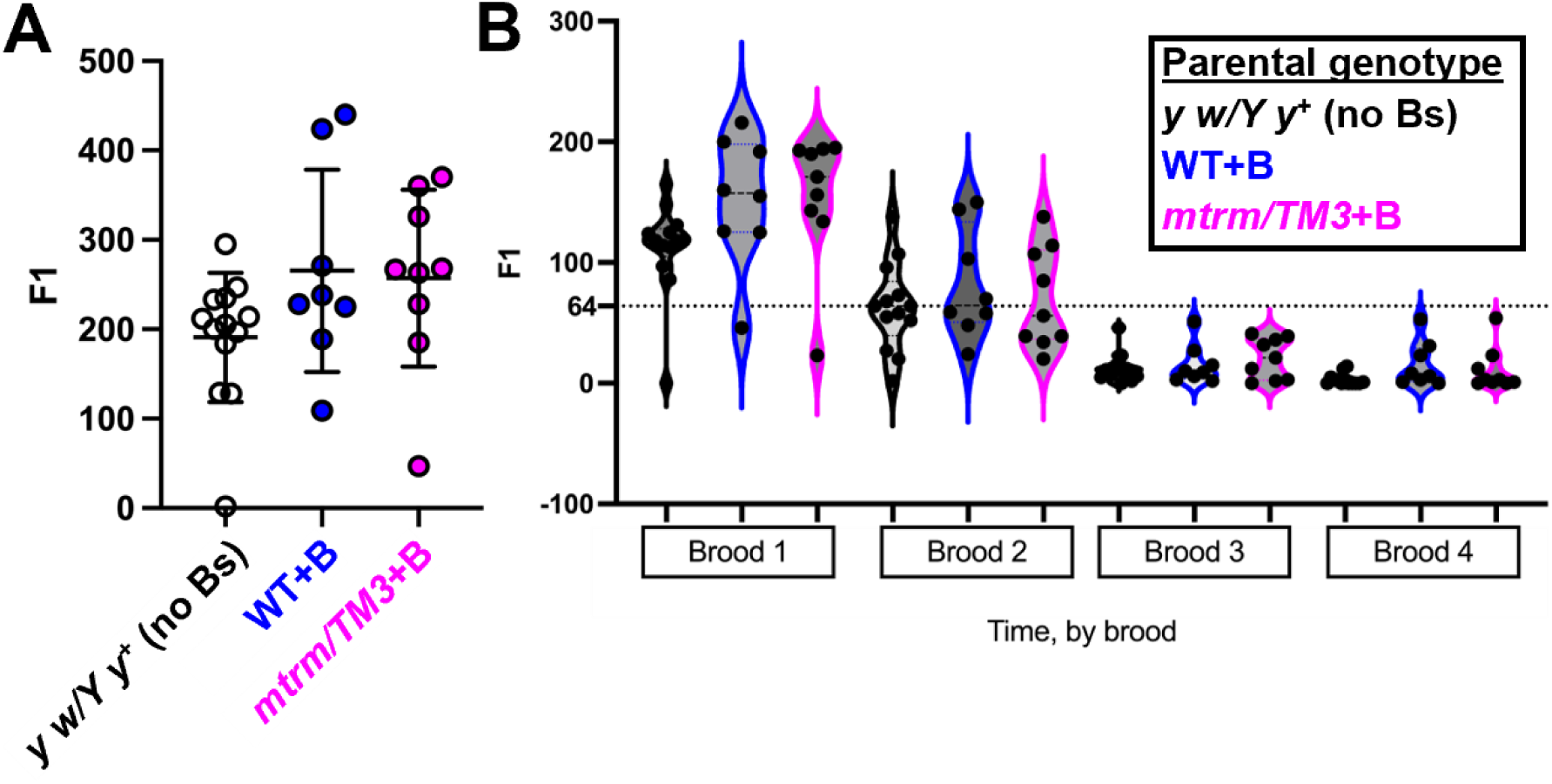
mtrm/TM3+B and WT+B males reveal heterogeneity in fertility. A) Total progeny (F1) sired by *y w/Yy^+^* males (mean ± standard deviation = 190.85 ± 72.18, n = 13), *WT+B* males (265.50 ± 113.23, n = 8), and *mtrm/TM3+B* males (257.11 ± 99.07, n = 9) from a sperm exhaustion assay. B) Violin plots of total progeny (F1) sired from a sperm exhaustion assay over time (by brood, see Methods). Black = progeny from *y w/Yy^+^* (no Bs) males, blue = progeny from *WT+B* males, magenta = progeny from *mtrm/TM3+B* males. Dashed line = median value for progeny from *y w/Yy^+^* (no Bs) brood 2. P-values calculated by One-Way ANOVA (Brown-Forsythe and Welch ANOVA tests) and Dunnett’s T3 multiple comparisons test; all calculated P values were not statistically significant (p > 0.05).

Over the span of the sperm exhaustion assay, on average, individual *mtrm/TM3+B* males and *WT+B* males showed no significant differences in total progeny over time when compared to each other and to *y w/Yy^+^* control males (Figure 1B, Supplemental Figure 1D-1F). Surprisingly, *mtrm/TM3+B* and *WT+B* males tended to sire more progeny on average in all broods compared to *y w/Yy^+^* control males (brood 1 = 155.44, 152.50, and 112.15, brood 2 = 70.22, 82.00, 63.38, brood 3 = 20.67, 15.38, 12.15, and brood 4 = 10.78, 15.63, and 3.15 respectively) (Figure 1B), allowing us to more confidently exclude the possibility that B chromosome drag occurs due to the selective loss of sperm with high B chromosome copy numbers. Though we are tempted to attribute this increase to the presence of B chromosomes, we cannot exclude the possibility that it is due to the presence of the marked fourth chromosome (*sv^spa-pol^*) or differences between the genetic background of the stocks.

### mtrm/TM3+B males exhibit heterogeneity in germline stem cell (GSC) hub morphologies

Since fertility differences cannot account for B chromosome drag in the *mtrm/TM3+B* genotype, we turned our attention to the beginning of spermatogenesis. We first examined the germline stem cells (GSCs) that are essential for the continued development of gametes and the transmission of genetic material to the next generation, as well as the hub cells that anchor GSCs to the apical tip of the testes. Through our observations using immunofluorescence, we found a heterogenous mix of GSC hub phenotypes among *mtrm/TM3+B* males that were completely absent in *WT+B* males (Figure 2A-D, Table 1). In *mtrm/TM3+B* testes, 25.8% had small or absent hubs based on Fasciclin III (Fas-III) staining (Figure 2B-C, Table 1), and 3.0% had tumorous and/or ectopic hubs (n = 66) (Figure 2D, Table 1). Despite these abnormal hub phenotypes, the average number of GSCs per testes appears to be unaffected as there was not a statistically significant difference between *WT+B* and *mtrm/TM3+B* males (Figure 2E).

**Figure 2:**
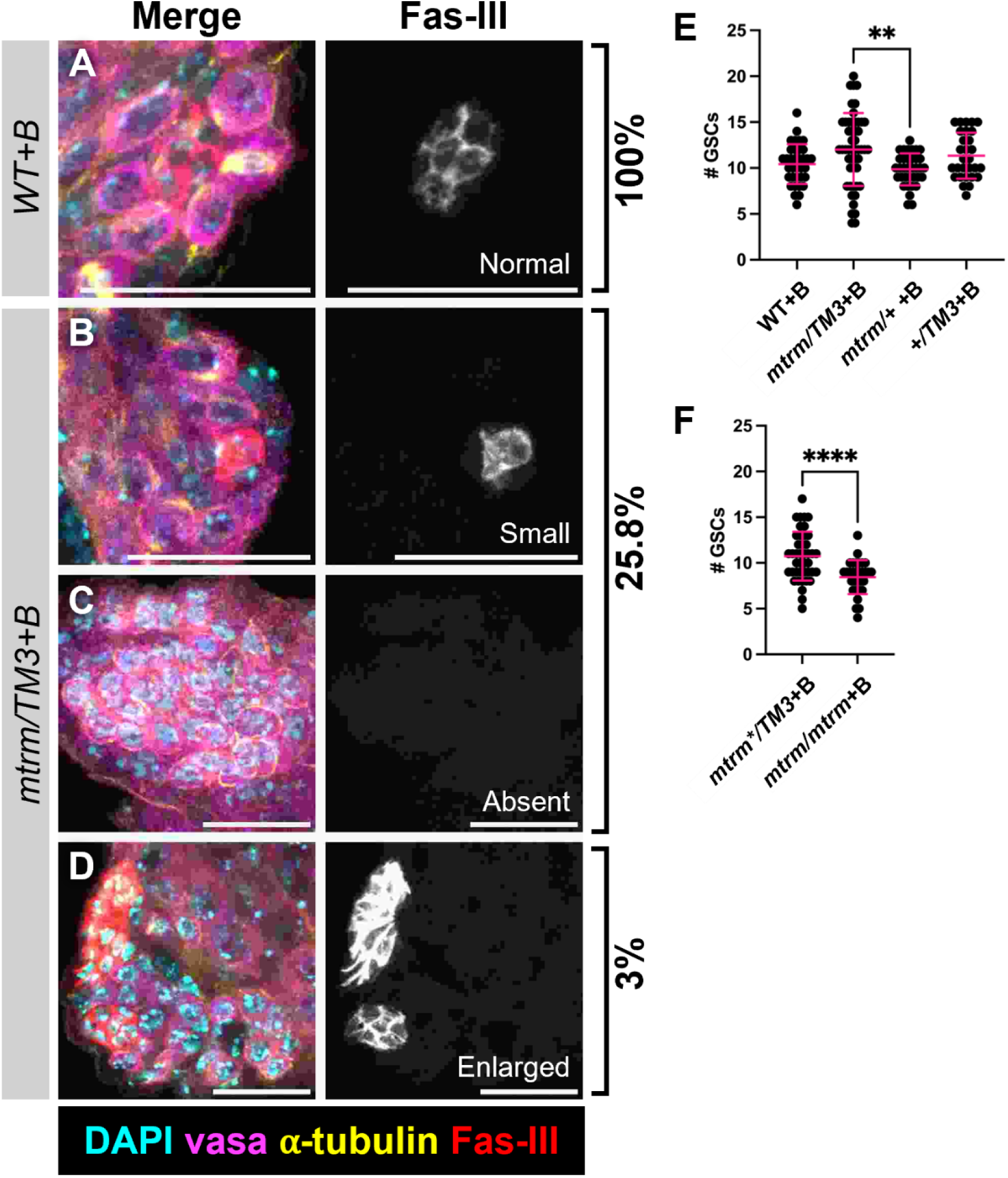
mtrm/TM3+B males display abnormal hub phenotypes. Representative immunofluorescence (IF) images of a typical *WT+B* male GSC niche (A), a *mtrm/TM3+B* male GSC niche with an abnormally small hub (B), a *mtrm/TM3+B* male GSC niche lacking a hub (C), and a *mtrm/TM3+B* male GSC niche with an ectopic hub (D). Cyan = DAPI, magenta = vasa, yellow = α-tubulin, red = Fas-III. All scale bars = 20 μm. E) Total number of GSCs per testis for *WT+B* (mean ± standard deviation = 10.43 ± 2.17, n = 35), *mtrm/TM3+B* (12.02 ± 3.98, n = 50), *mtrm/+ +B* outcrossed males (9.86 ± 1.73, n = 35), and *TM3/+ +B* outcrossed males (11.34 ± 2.50, n = 29). P-values calculated by One-Way ANOVA (Brown-Forsythe and Welch ANOVA tests) and Dunnett’s T3 multiple comparisons test. F) Total number of GSCs per testis for *matrimony* transheterozygous mutant null sibling control males (mean ± standard deviation = 10.74 ± 2.67, n = 42) and *matrimony* transheterozygous mutant null males (8.45 ± 1.86, n = 31). P-values calculated by unpaired t-test with Welch’s correction. ns = p > 0.05, ** = p ≤ 0.01, **** = p ≤ 0.0001.

**Table 1:** Qualitative morphological assessment of the somatic hub of the male GSC niche.

| Genotype (full) | total hubs<br>quantified | # of testes<br>with small or<br>no hubs | # of testes<br>with extra or<br>tumorous<br>hubs | % small<br>or no<br>hubs | % extra or<br>tumorous<br>hubs |
| --- | --- | --- | --- | --- | --- |
| *y w/Y,y<sup>+</sup>;sv<sup>spa-pol</sup> +B<sup>a</sup>* | 44 | 0 | 0 | 0.0 | 0.0 |
| *y w/Y,y<sup>+</sup>;mtrm<sup>126</sup>/TM3, Sb Ser;sv<sup>spa-pol</sup> +B<sup>b</sup>* | 66 | 17 | 2 | 25.8 | 3.0 |
| *y w/Y,y<sup>+</sup>;TM3, Sb Ser/+;sv<sup>spa-pol</sup> +B<sup>a</sup>* | 29 | 0 | 0 | 0.0 | 0.0 |
| *y w/Y,y<sup>+</sup>;mtrm<sup>126</sup>/+;sv<sup>spa-pol</sup> +B<sup>b</sup>* | 35 | 0 | 0 | 0.0 | 0.0 |
| *y w/Y,y<sup>+</sup>; mtrm<sup>126</sup> or Df(3L)/TM3,Sb Ser; sv<sup>spa-pol</sup> +B<sup>c</sup>* | 43 | 0 | 0 | 0.0 | 0.0 |
| *y w/Y,y<sup>+</sup>; mtrm<sup>126</sup>/Df(3L)66C-T2-10, nanos-GAL4; sv<sup>spa-pol</sup> +B<sup>c</sup>* | 35 | 0 | 0 | 0.0 | 0.0 |

<sup>a</sup>2-4 days old, <sup>b</sup>2-3 days old, <sup>c</sup>1-3 days old

The lack of abnormal hub morphologies in *WT+B* males allowed us to eliminate the B chromosomes as the source of this phenotype, therefore we wanted to determine whether *mtrm* or the *TM3* balancer chromosome was responsible for hub disruption. To test this, we outcrossed the *mtrm/TM3+B* stock to wild-type and characterized hub morphology and average number of GSCs in *mtrm/+ +B* and *+/TM3+B* males (see Supplemental Table 1 for outcross genotypes). Neither of the genotypes from the outcross resulted in abnormal hubs, and the average number of GSCs was not statistically significantly different from *WT+B* (Figure 2E, Table 1), leading us to initially conclude that the combination of *mtrm* and *TM3* is required for hub disruption. We attempted to validate this conclusion by outcrossing the *mtrm/TM3+B* stock to a stock carrying a different mutant allele of *mtrm* over the *TM3* balancer to produce *mtrm*/TM3+B* males (we were unable to distinguish which allele of *mtrm* the progeny inherited, therefore this genoype is designated with an * instead of a specific allele). To our surprise, we observed no abnormal hub phenotypes and no statistically significant difference in average number of GSCs (Figure 2F, Table 1), indicating that the abnormal hub phenotype is not a direct result of *mtrm*, *TM3*, or a combination of the two; rather, it is likely the product of an uncharacterized genetic polymorphism that exists exclusively in the *mtrm/TM3+B* stock.

Though we found that the average number of GSCs in *mtrm/+ +B* testes was similar to *WT+B*, we were intrigued by the result that the average was statistically significantly lower than *mtrm/TM3+B* (Figure 2E). Previous studies showed that Matrimony protein is not present in adult male testes (Xiang et al. 2007), however, Fly Cell Atlas reports that *matrimony* transcript is detected at low levels in male gonadal cells (Li et al. 2022). To further test if the loss of *matrimony* affects normal GSC morphology, we collected transheterozygous *mtrm/mtrm+B* males that were the siblings of the *mtrm*/TM3+B* males described above (Supplemental Table 1). Transheterozygous *mtrm/mtrm+B* mutant flies had no abnormal hubs compared to heterozygous sibling controls (Table 1), however they did have significantly fewer GSCs on average compared to their sibling controls (Figure 2F). The trends in our data are currently consistent with two hypotheses: either the *TM3* balancer chromosome promotes a modest increase in the average number of GSCs, or *mtrm* has a minor but detectable role in promoting GSC development. Additional studies will be necessary to distinguish between these possibilities.

### B chromosomes are symmetrically segregated during male germline stem cell (GSC) divisions

The only genotype that exhibited abnormal hub morphology was the *mtrm/TM3+B* stock that is capable of B chromosome drag. We reasoned that abnormal hubs may be correlated with irregular GSC divisions, which could be the source of B chromosome drag. We first measured the mitotic index of GSCs to determine the proportion that were actively undergoing division (identified by α-tubulin and DAPI of GSCs in metaphase, anaphase, or telophase). If mitotic indexes are higher in *mtrm/TM3+B* males, it may be from frequent activation of the spindle assembly checkpoint that could increase the frequency of lagging B chromosomes, which then become physically separated from the genome and are eventually lost. we found no statistically significant difference in the percent of dividing GSCs (Supplemental Figure 2B and 2D), indicating the proportion of the cell cycle that is mitosis is roughly the same across genotypes.

Though mitotic timing is the same across genotypes, the pattern of B chromosome segregation in these mitoses may vary. The GSC division is inherently asymmetric because it produces one GSC and one gonialblast (GB) cell, and we were curious if the B chromosomes segregated symmetrically or asymmetrically. To test this, we measured the total *AAGAT* FISH signal, which is enriched on the B chromosomes and chromosome 4 (Hanlon et al. 2018), as a proxy for the ratio of B chromosomes distributed between a GSC and its recent GB decedent (Figure 3A-C). Since chromosome 4 segregation in males is not significantly affected by B chromosomes (Bauerly et al. 2014) or the loss of *mtrm* (Harris et al. 2003), we presume that comparing *AAGAT* fluorescence intensity ratios would be minimally affected by chromosome 4 signal and would represent any differences in B chromosome abundance.

**Figure 3:**
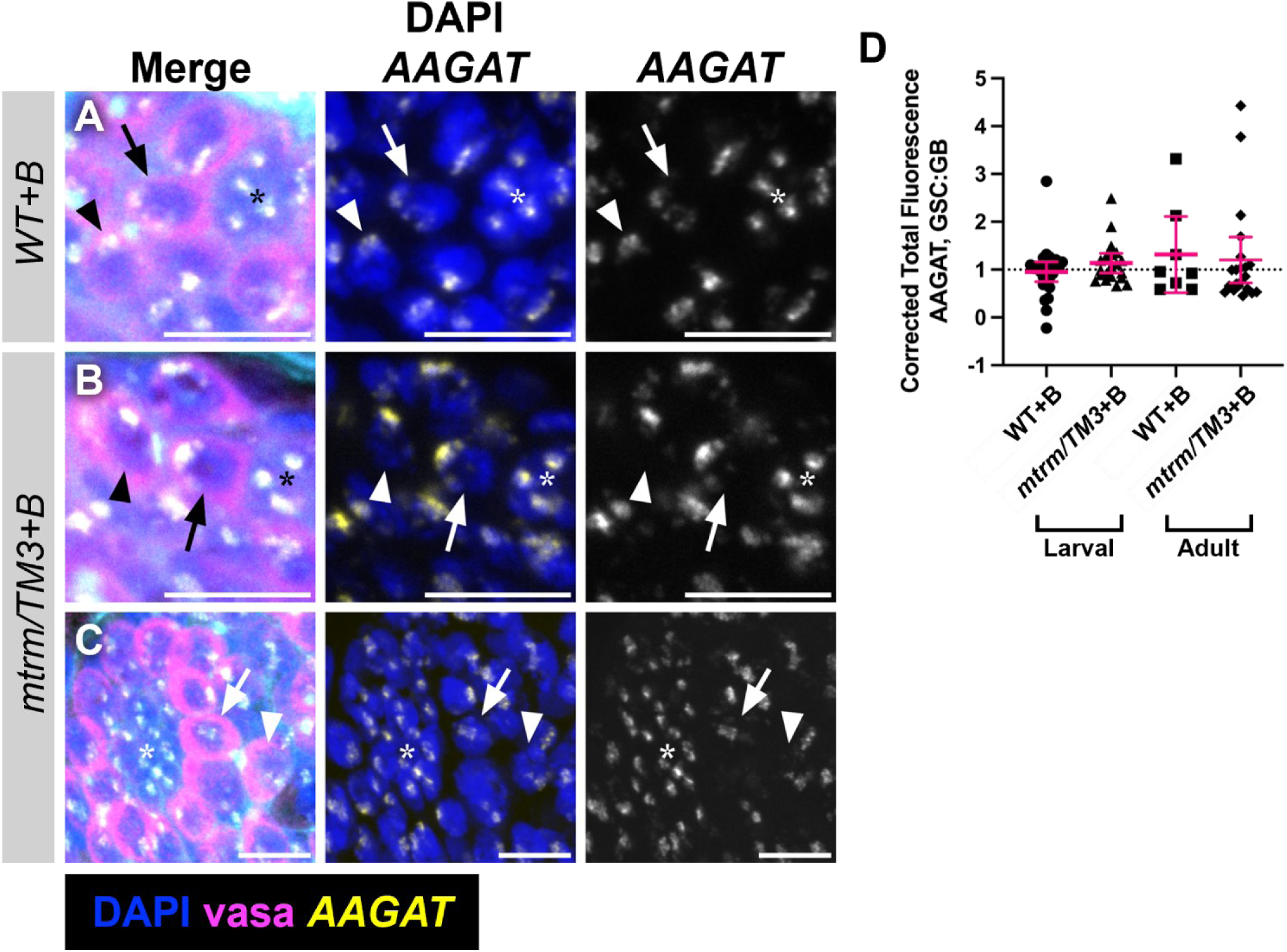
B chromosomes are symmetrically segregated during GSC divisions. Representative IF+FISH images of GSC:GB pairs of adult *WT+Bs* adult males (A), *mtrm/TM3+Bs* adult males (B), and *mtrm/TM3+Bs* larval males (C). All scale bars = 10 μm. Blue = DAPI, magenta = vasa, cyan = α-spectrin, yellow = *AAGAT.* D) Corrected total fluorescence (CTF) of *AAGAT* ratio of GSC value to GB value (GSC:GB) for *WT+B* adult males with significant outlier removed (p < 0.05 calculated by Grubb’s test) (mean ± standard deviation = 0.95 ± 0.52, n = 27), *mtrm/TM3+B* adult males (1.14 ± 0.44, n = 20), *WT+B* larval males (1.32 ± 0.95, n = 8), and *mtrm/TM3+B* larval males (1.20 ± 1.05, n = 21). P-values calculated by One-Way ANOVA (Brown-Forsythe and Welch ANOVA tests) and Dunnett’s T3 multiple comparisons test. ns = p > 0.05. E) Model of our current understanding of where paternal B chromosome drag may be occurring.

The first sperm generated by males undergo meiosis during larval development and are the first sperm utilized in our B chromosome transmission assays and the sperm exhaustion assays described above. Therefore, we assayed *AAGAT* signal in GSC:GB pairs of third instar larval testes as well as young adult testes. We expected to see values less than 1 if there was a trend to reduce the number of B chromosomes distributed to GBs, however we found no statistically significant differences between the corrected total fluorescence (CTF) of *AAGAT* GSC:GB ratio of *mtrm/TM3+B* compared to *WT+B* males for both third instar larval and adult males, with or without the removal of the outlier value of *WT+B* adult males (significant outlier of p < 0.05 calculated by Grubb’s test) (Figure 3D, Supplemental Figure 3). Together with our previous findings, we conclude that the mechanism for B chromosome drag most likely occurs beyond the GSC niche.

## DISCUSSION

Selfish genetic elements achieve non-Mendelian transmission by generating or exploiting asymmetries during sexual reproduction. Unlike female meiosis where only one of the four meiotic products are retained, all four products of male meiosis go on to become sperm. For drive to occur during male meiosis, a selfish genetic element must either generate asymmetry, (e.g., killing its competition) or take advantage of an asymmetry that naturally exists, such as the GSC:GB division. In the case of *mtrm/TM3+B* males, to produce the opposite effect (drag), sperm with higher numbers of B chromosomes would have to be eliminated so that sperm with lower numbers would be in abundance, or B chromosomes would have to be unevenly distributed during the asymmetric GSC:GB divisions. Our study initiates the investigation into the mechanism behind B chromosome drag and eliminates sperm inviability and unequal distribution of B chromosomes during the GSC:GB division as possible sources of drag.

Despite finding no statistically significant differences between presumed B chromosome inheritance patterns after GSC divisions (Figure 3D), the slightly elevated average GSC:GB ratio in larval testes and adult *mtrm/TM3+B* testes suggests that there may be a subtle bias towards B chromosome inheritance into the GSC in larval testes that continues into adulthood for *mtrm/TM3+B* males but not *WT+B* males. If paternal B chromosome counts were based on cells derived from GSCs accumulated B chromosomes, then it may appear that B chromosome transmission is reduced when using B chromosome copy numbers obtained from initial/early progeny. We disfavor this possibility because the B chromosome transmission assays use a young virgin male that is mated with a female for one day before he is removed, meaning that GSCs may not have undergone enough divisions to allow it to accumulate B chromosomes.

Based on our cumulative results, we suggest that the predominant force(s) impacting B chromosome drag may occur outside of the GSC niche. Immediately downstream from the GSC niche is where cells divide mitotically to reach 16-cell clusters. These cells then enter into the meiotic program that promotes chromosome territory formation, homolog pairing, and meiotic chromosome segregation. After meiosis is finished, each of the 64-cell bundles undergo a series of transformations to development into mature sperm, including the programmed elimination of their mitochondria and the exchange of nucleosomes for protamines. Future studies that aim to identify the source of the B chromosome drag will likely apply a similar process of elimination that we employed here to these mitotic, meiotic, and developmental stages. Altogether, we are hopeful that this emerging model system of B chromosome dynamics will reveal novel points of asymmetry during spermatogenesis and mechanisms that allow the germline to promote disparate chromosome behavior.

## ACKNOWLEDGEMENTS

We would like to thank the members of the Hanlon lab and Dr. Barbara Mellone and the Mellone lab members for their advice and feedback on this research. Thank you to Dr. Mayu Inaba and lab members for the Fasciclin-III (Fas-III) antibody. The Leica SP8 Spectral Confocal microscope housed within the Advanced Microscopy Facility at the University of Connecticut was acquired with NIH grant S10OD016435 awarded to Akiko Nishiyama.

---

**Supplemental Figure 1:**
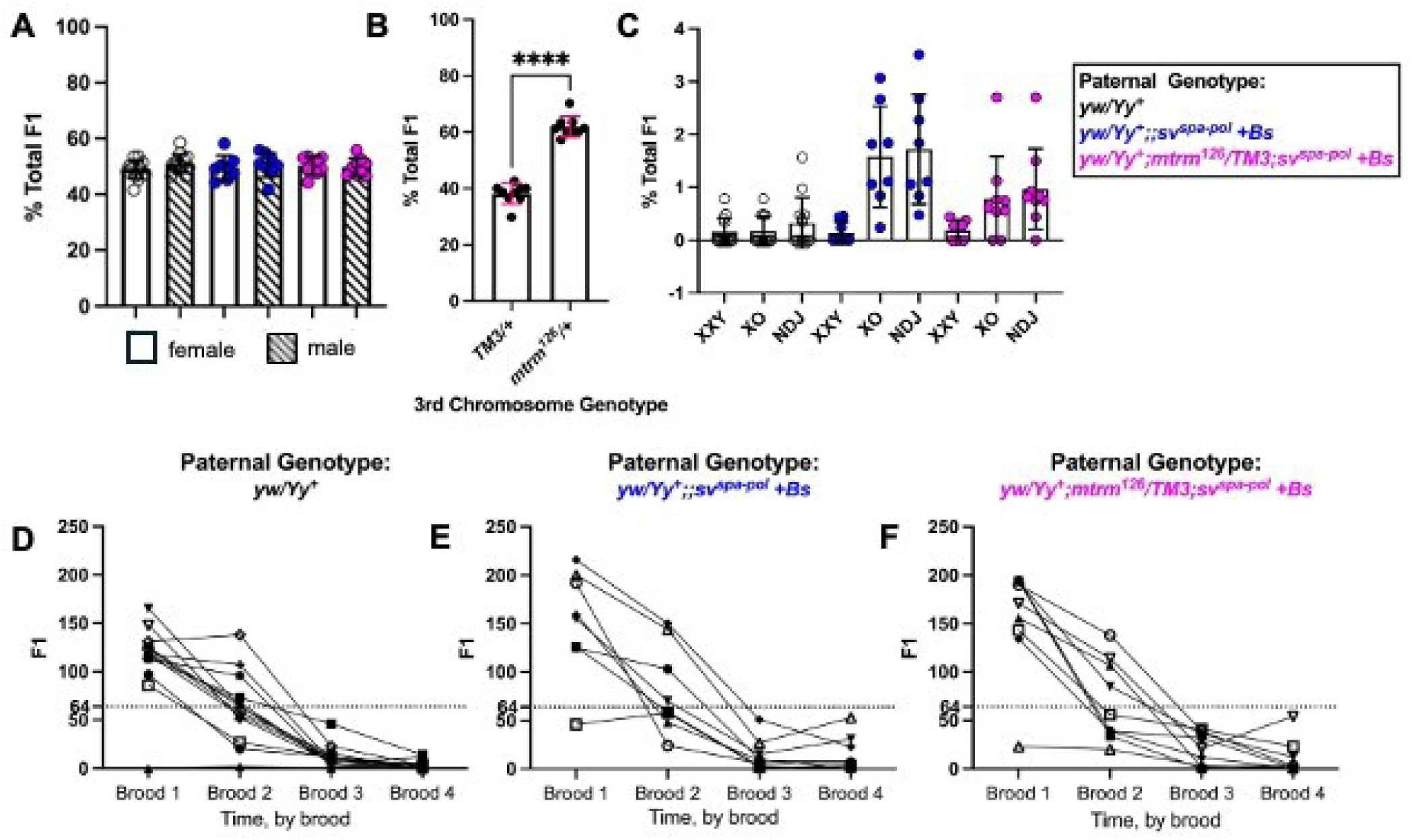
Sperm exhaustion assay details. A) Percent sex of total progeny (F1) by paternal genotype. Empty bars = female, bars with angled stripes = male. Female F1 and male F1 respectively from *y w/Yy^+^* males (mean ± standard deviation = 48.96 ± 3.33, and 51.04 ± 3.33, n =13), *WT+B* males, (49.55 ± 4.34 and 50.45 ± 4.34, n = 8), and *mtrm/TM3+B* males (50.25 ± 3.10 and 49.75 ± 3.10, n = 9). P-values calculated by One-Way ANOVA (Brown-Forsythe and Welch ANOVA tests) and Dunnett’s T3 multiple comparisons test. B) Percent of total progeny (F1) inheriting paternal third chromosome, *mtrm* or *TM3* respectively (mean ± standard deviation = 61.88 ± 3.64 and 38.12 ± 3.64, n = 9). P-values calculated by unpaired t-test with Welch’s correction. C) Percent of total progeny (F1) with sex chromosome non-disjunction event(s) by paternal genotype. XXY = *y w/y w/Yy^+^* female, XO = *y w/O* male, and NDJ = XXY+XO respectively for *y w/Yy^+^* males (mean ± standard deviation = 0.15 ± 0.26, 0.17 ± 0.28, and 0.32 ± 0.48, n =13), *WT+B* males, (0.14 ± 0.21, 1.58 ± 0.95, and 1.72 ± 1.04, n = 8), and *mtrm/TM3+B* males (0.19 ± 0.18, 0.78 ± 0.80, and 0.97 ± 0.76, n = 9). Black = progeny from *y w/Yy^+^* males, blue = progeny from *WT+B* males, magenta = progeny from *mtrm/TM3+B* males. P-values calculated by One-Way ANOVA (Brown-Forsythe and Welch ANOVA tests) and Dunnett’s T3 multiple comparisons test. ns = p > 0.05, ** = p ≤ 0.01, **** = p ≤ 0.0001. F1 production over time (by brood, see Methods) for individual *y w/Yy^+^* (D), *WT+B* (E), and *mtrm/TM3+B* (F) males. Dashed line = median value for *y w/Yy^+^* brood 2 progeny. No statistics calculated for D-F.

**Supplemental Figure 2:**
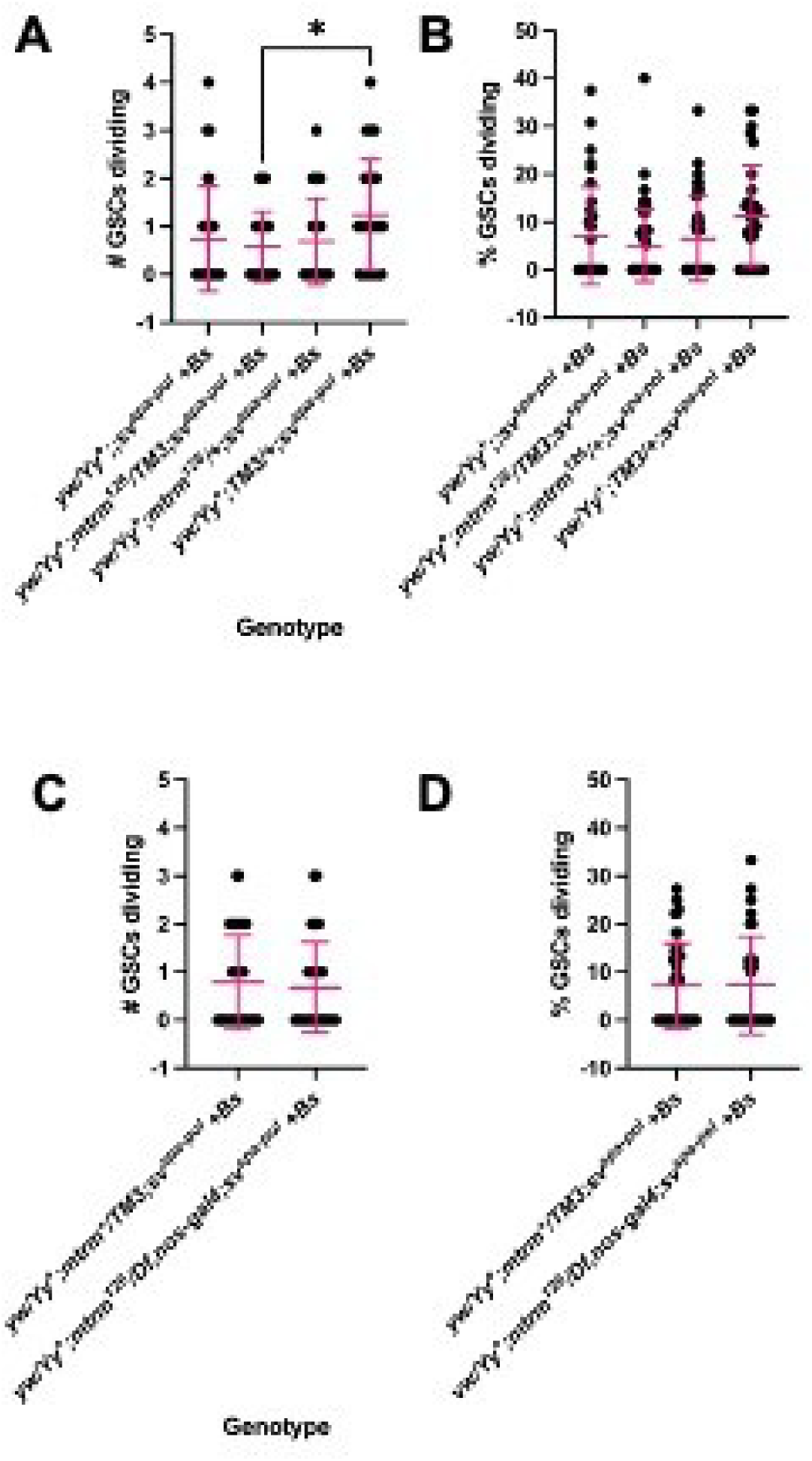
Actively dividing GSCs. Total number (A, C) and percentage (B, D) of GSCs per testis actively dividing for *WT+B* males (Total number: mean ± standard deviation = 0.74 ± 1.09, Percentage: 7.10 ± 10.16, n = 35), *mtrm/TM3+B* males (Total number: 0.58 ± 0.73, Percentage: 5.09 ± 7.56, n = 50)*, mtrm/+ +B* outcross males (Total number: 0.69 ± 0.90, Percentage: 6.47 ± 8.78, n = 35), and *TM3/+ +B* outcross males (Total number: 1.24 ± 1.15, Percentage: 11.13 ± 10.70, n = 29) P-values calculated by One-Way ANOVA (Brown-Forsythe and Welch ANOVA tests) and Dunnett’s T3 multiple comparisons test (A-B) and for *matrimony* transheterozygous mutant null sibling control males (Total number: 0.79 ± 0.98, Percentage: 7.16 ± 8.86, n = 42) and *matrimony* transheterozygous mutant null males (Total number: 0.68 ± 0.94, Percentage: 7.25 ± 10.19, n = 30) (C-D) P-values calculated by unpaired t-test with Welch’s correction. ns = p > 0.05.

**Supplemental Figure 3:**
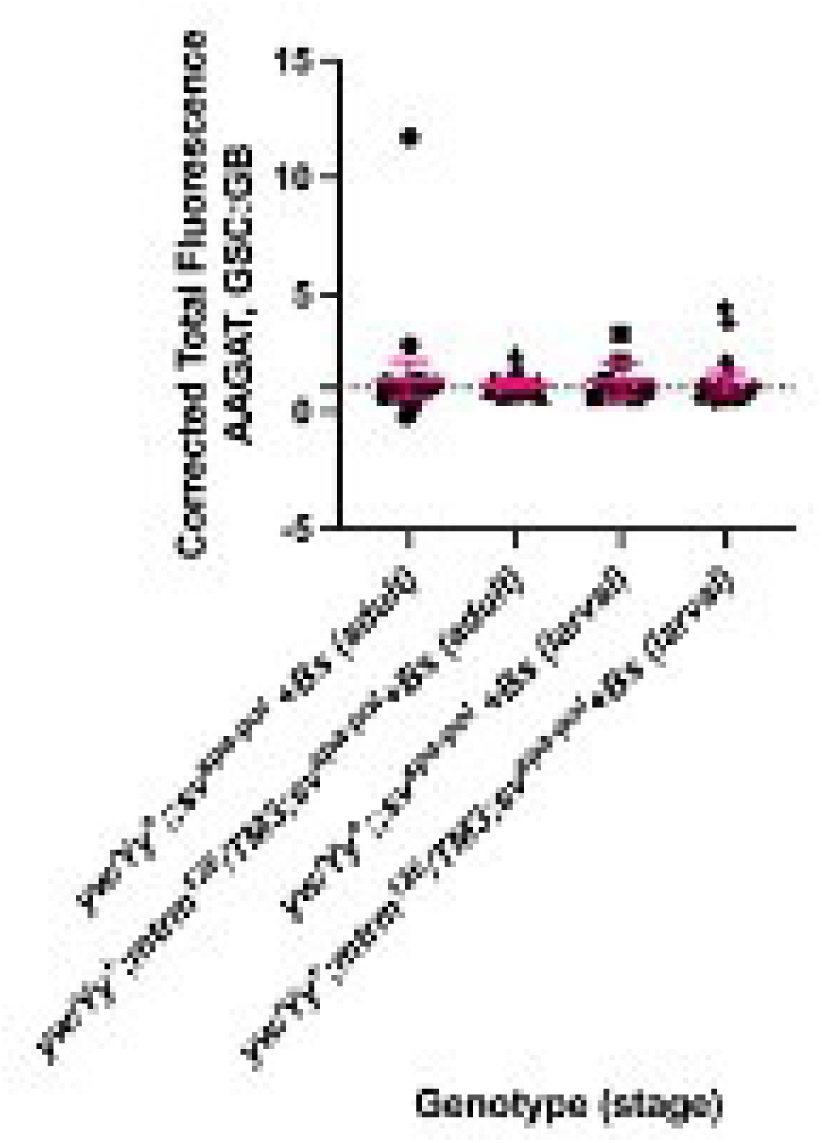
CTF of GSC:GB ratios including an outlier. Corrected total fluorescence (CTF) of *AAGAT* ratio of GSC value to GB value (GSC:GB) for *WT+B* adult males including the significant outlier (p < 0.05 calculated by Grubb’s test) (mean ± standard deviation = 1.34 ± 2.09, n = 28), *mtrm/TM3+B* adult males (1.14 ± 0.44, n = 20), *WT+B* larval males (1.32 ± 0.95, n = 8), and *mtrm/TM3+B* larval males (1.20 ± 1.05, n = 21). P-values calculated by One-Way ANOVA (Brown-Forsythe and Welch ANOVA tests) and Dunnett’s T3 multiple comparisons test. ns = p > 0.05.

**Supplemental Table 1.** Explanation of stocks and genotypes used in this study.

| Short name | Genotype | Notes | Carries Bs? |
| --- | --- | --- | --- |
| *y,w/Y,y<sup>+</sup>* (no Bs) | *y,w/Y,y<sup>+</sup>* | *y,w/Y,y<sup>+</sup>* | no |
| *WT* | *y,w/Y,y<sup>+</sup>;sv<sup>spa-pol</sup>* | DSLH5, Genetic wildtype | no |
| *WT+B* | *y,w/Y,y<sup>+</sup>;sv<sup>spa-pol</sup> +Bs* | -- | yes |
| *mtrm/TM3+B* | *y,w/Y,y<sup>+</sup>;mtrm<sup>126</sup>/TM3,Sb,Ser;sv<sup>spa-pol</sup> +Bs* | DSLH17 v2 | yes |
| *mtrm/+ +B* | *y,w/Y,y<sup>+</sup>;mtrm<sup>126</sup>/+;sv<sup>spa-pol</sup> +Bs* | Outcross males | yes |
| *+/TM3+B* | *y,w/Y,y<sup>+</sup>;TM3,Sb,Ser/+;sv<sup>spa-pol</sup> +Bs* | Outcross males | yes |
| *matrimony deficiency stock* | *y,w/Y,y<sup>+</sup>; Df(3L)66C-T2-10, nanos-GAL4/TM3, Sb Ser; sv<sup>spa-pol</sup> +Bs* | DSLH18 | yes |
| *mtrm/mtrm+B* | *y,w/Y,y<sup>+</sup>; mtrm<sup>126</sup>/Df(3L)66C-T2-10, nanos-GAL4; sv<sup>spa-pol</sup> +Bs* | *mtrm* transheterozygous mutant | yes |
| *mtrm\*/TM3+B* | *y,w/Y,y<sup>+</sup>; mtrm<sup>126</sup>/TM3,Sb,Ser; sv<sup>spa-pol</sup> +Bs* | Genotypes are indistinguishable | yes |
|  | *y,w/Y,y<sup>+</sup>; Df(3L)66C-T2-10, nanos-GAL4/TM3,Sb,Ser; sv<sup>spa-pol</sup> +Bs* |  | yes |

